# Effects of dose-response relationships on cellular information transmission

**DOI:** 10.64898/2026.10.26.761777

**Authors:** Steven S. Andrews, H. Steven Wiley, Roger Brent, Herbert M. Sauro

**Affiliations:** Department of Bioengineering, University of Washington, Seattle, Washington, USA; Environmental Molecular Sciences Laboratory, Pacific Northwest National Laboratory, Richland, Washington, USA; Division of Basic Sciences, Fred Hutchinson Cancer Research Center, Seattle, Washington, USA

## Abstract

Cell signaling systems can transmit information with high precision, despite relying on complex reaction networks and inherently noisy biochemical reactions. To better understand how they achieve this precision, this work focuses on how cells can tune their quantitative steady-state dose-response relationships to improve signaling fidelity. Using a combination of analytic and numerical approaches, we quantify the maximum information that can be transmitted in single-step and multi-step signaling systems under several biologically realistic noise distributions: Gaussian, gamma, constant coefficient of variation, and binomial. We find that signaling systems transmit more information if their dose-response curves have low baselines, large dynamic ranges, and curve shapes that are well matched to the natural distribution of input signals. For multi-step systems, linear transfer functions typically improve signaling fidelity for all noise levels, consistent with previous work on “dose-response alignment”. We also find that switchlike transfer functions transmit a maximum of one bit and are robust to low levels of noise, but fail rapidly with high levels of noise. Simulations of toy signaling systems corroborate these results. Together, these findings provide insight into some cellular design principles that have arisen through evolution and that are widely observed in nature.

**Author summary:** Cell signaling systems transmit information about extracellular conditions into the cell, where it can be processed and acted upon. At the steady state, these system responses can be described by dose-response curves that relate system output to input. Because the shapes of these curves can presumably be tuned through evolution, this raises the question of what shape dose-response curves maximize the accuracy of signal transmission. Using a combination of analytical theory and numerical calculations, we find that signaling systems transmit the most information when their dose-response curves are matched to the natural distribution of input signals. In signaling pathways composed of multiple sequential steps, information transmission is typically maximized when downstream signaling steps respond linearly to the outputs of upstream steps. In principle, cells could also use switchlike responses, in which signals are either on or off, but we show that these approaches fail rapidly with high noise levels. These predicted relationships agree with simulation results and are broadly consistent with observations of natural signaling systems.

## Introduction

Signal transduction pathways transmit information about extracellular conditions into cells using networks of biochemical reactions. The cells then use the information to make decisions that affect their behaviors. For example, a cell may make decisions about which way to swim [1], or whether it should grow, divide, differentiate, mate, or undergo apoptosis [2, 3].

The ability to make correct decisions affects an organism’s fitness, so one would reasonably expect that many of these signaling pathways would have evolved to transmit information with good precision. Indeed, high-precision signaling is widely observed. For example, receptors in human visual systems signal sufficiently well that they can count single photons [4], *Bombyx* moth olfactory systems can detect single odorant molecules [5], *Dictyostelium discoideum* cells can detect differences of only about 10 molecules between the front and back of the cell [6], HeLa cells can reliably discriminate between at least 5 different insulin growth factor doses [7], and morphogen signaling in developing embryos is sufficiently precise to define unique identities for individual cells [8, 9]. In addition, we recently quantified signaling precision in the yeast cell pheromone response signaling system and found that it transmits signals sufficiently precisely to convey over 3 bits of information in an individual static signal [10]. This is sufficient for a cell to reliably distinguish between about 2^3^ = 8 different extracellular pheromone concentrations.

These findings of precise signaling lead to the question of how cells maintain precision despite inherent biochemical noise that arises from the small copy numbers of intracellular proteins and transcripts [11].

This noise is substantial. For example, Poissonian noise for a signaling protein that exists with 100 copies would imply a variation of 10 molecules, which is 10%. Furthermore, noise that arises over multiple pathway steps is cumulative, and noise can also be amplified, such as during the transcription of small numbers of mRNA transcripts to large numbers of proteins [12, 13]. Additionally, intracellular spatial organization can increase sensitivity to events that occur in small volumes of space, where there tends to be more copy number variation [11].

Previous research on how cells achieve accurate signaling has revealed many possible mechanisms [14]. (1) Signaling noise can be reduced by using more signaling molecules [15]. (2) Faster activation-deactivation cycles of signaling molecules in upstream portions of signaling pathways, and slower cycles in downstream portions, allow better signal averaging [16]. (3) Some signaling network topologies naturally compensate for noise, of which feedforward loops [17–19] and negative feedback loops [20–23] can be particularly effective. (4) Functional redundancy, in which the signal is carried by multiple pathways, can mitigate the effects of noise introduced at any one point [24, 25]. (5) Protein dimerization and oligomerization can buffer protein concentrations and thus reduce the effects of stochastic variability [26–28]. (6) Dynamic signaling, in which the signal is measured over time, can improve information transfer both through averaging results over multiple measurements and by enabling removal of noise that arises from cell-to-cell variation [29]. Finally, vice versa (7), cell-to-cell variation can also benefit information transfer to a cell population because different cells can have peak sensitivities at different input concentrations [30–32]. These mechanisms are not mutually exclusive but may act together to yield combined benefits. It is also worth noting that many of these mechanisms have costs associated with them, including particularly the metabolic costs of synthesizing more components such as proteins and mRNAs.

Here, we investigate yet another mechanism that cells may use to improve signaling precision: cells may have tuned their steady-state dose-response curves between signaling system inputs and outputs, through evolution, to improve signal transmission. This idea was first proposed by Laughlin [33], who showed that blowfly visual neuron contrast-response functions closely matched the contrast levels in natural scenes. This idea also builds on observations that many cell signaling systems, including the insulin [34], acetylcholine [35], thyroid stimulating hormone [36], angiotensin II [37], epidermal growth factor (EGF) [38–41], Wnt [41], TGFβ [41], and yeast pheromone response [42, 43] signaling systems all exhibit nearly linear signal transmission at one or more stages within their signaling pathways. Some of us dubbed this phenomenon “dose-response alignment” [43, 44], or DoRA, to reflect the fact that multiple steps of a signaling pathway exhibit dose-response functions that nearly align with each other [45]. We suggested previously that linear signaling may improve information transmission because relative signal strengths are maintained throughout the signaling pathway, rather than being rescaled to higher or lower values [43–47].

We revisit this suggestion here, rigorously quantifying information transmission for signaling over single and multiple pathway steps in model signaling systems. Here, a “step” is any portion of a signaling pathway that has well-defined inputs and outputs; it could be as simple as a single protein that can be either phosphorylated or dephosphorylated, or as complex as an entire signaling pathway. For signaling over one step, we present information transmission equations for several biologically realistic noise distributions.

These equations show that signaling pathways transmit more information if their dose-response curves have low basal responses, high dynamic ranges, and are adapted to the natural signal distribution. For signaling over two steps, we confirm prior suggestions that linear signaling can improve information transmission [43–47]. However, we also show that linearity is only strictly optimal if each pathway step has the same noise distribution and, even then, it is only one of a class of optimal distributions. The same principles apply to longer pathways as well. Switchlike signaling provides an interesting contrast to the more common graded dose-response curves and is remarkably robust to low levels of noise. However, it fails rapidly with high levels of noise. Simulation results confirm these theoretical predictions.

### Information theory background

Information theory was initially developed for investigating telecommunications [48], but is just as valid for investigating any other system that transmits information. While it is mathematically complicated, it is used because its measure of information, called the Shannon entropy, is the only mathematical function that obeys several intuitive rules about how information ought to behave [48, 49] (e.g. predictable outcomes have zero information content and independent information amounts are additive). This unique suitability has led it to be applied increasingly to cell signaling over the past few decades [10, 24, 50–55]. Here, we summarize some particularly relevant parts of the theory and define our terms, while directing readers to books and reviews [48, 56–62] for more general background.

For signaling over one step, we consider signaling from an input, S, to an output, R, through a noisy signaling system, or channel (Fig. 1A). For concreteness, it helps to think about signaling within an individual cell, where a typical signal value might be an extracellular ligand concentration, a typical response value could be a protein activity level or a gene expression rate, and the noise would arise from within the cell’s signaling pathway. This type of noise, called intrinsic noise [10, 63–66], limits the amount that a cell is able to learn about its environment. However, the results given below are sufficiently general that they also apply to multicellular signaling, molecular communication [67, 68], and other situations. They also apply to extrinsic noise, including particularly variation between different individual cells.

**Fig 1.**
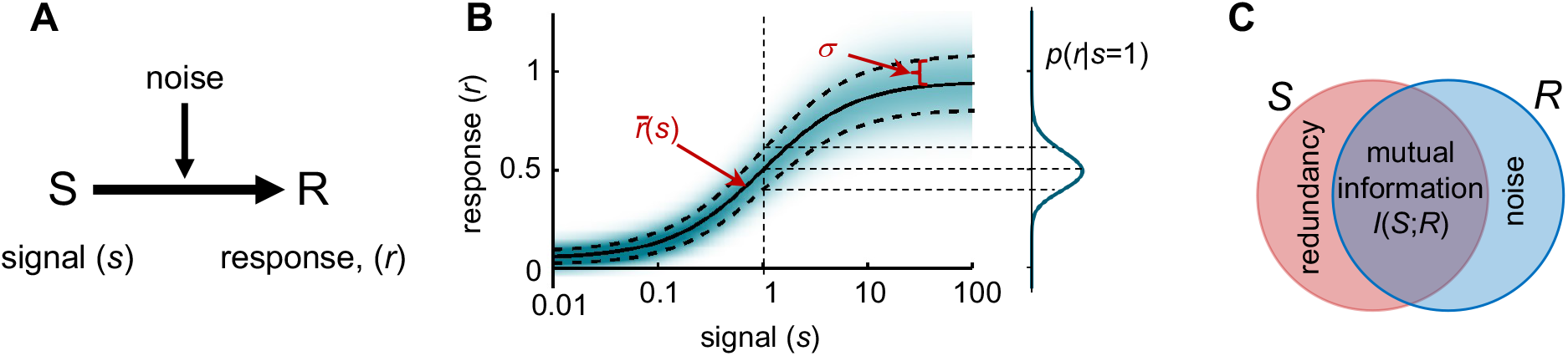
Information theory background. (A) Diagram of signaling from signal, S, to response, R, with added noise. (B) Signal-response-variation (SRV) plot for steady-state signaling, illustrating the conditional response distribution with shading, the mean response (the dose-response curve) with a solid black line, and the response standard deviation with thick dashed lines. The curve on the right illustrates the response probability distribution that would be observed for a signal value of 1. In this illustration, the dose-response function is a Hill function with *A* = 0.9, *B* = 0.05, *E* = 1, and *N* = 1 and the noise is gamma distributed with coefficient *n*_Γ_ = 0.14 (see main text). (C) Venn diagram of signal information in red and response information in blue. The overlapping region represents the mutual information. Additional information in the response represents information about the noise and additional information in the signal represents signal values that produce redundant responses.

We focus on the steady state, so this system is fully mathematically defined by two functions: the signal distribution, *p*(*s*), which represents the likelihood of any particular signal value being sent, and the conditional response distribution, *p*(*r* |*s*), which represents the probability of observing response *r* for the given signal value *s* [24, 69]. *The main part of Fig. 1B, which we call a signal-response-variation (SRV) plot [10], illustrates p*(*r* |*s*) with shading and the graph on the right of the figure illustrates *p*(*r* |*s*) for the case where *s* = 1.

The solid black line in the figure represents the dose-response function, which is the mean response as a function of the signal,

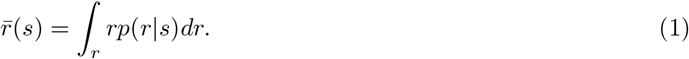

For convenience, we assume that the dose-response function has a finite range, from 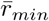 to 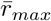, and increases monotonically and in a graded fashion. These restrictions are not necessary but are often valid and simplify the mathematics. Our figures and numerical work represent the dose-response function with a Hill function, given by

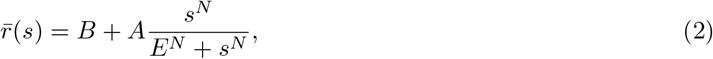

where *B* is the baseline, *A* is the amplitude, *E* is signal value that yields half-response and abbreviated as EC_50_, and *N* is the Hill cooperativity [46, 70]; in this case, 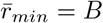 and 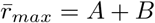. The Hill function generalizes the Michaelis-Menten equation by adding baseline and cooperativity parameters, and also shifts the focus from enzyme-controlled reaction rates to general dose-response relationships. Note that the figure follows convention by using a logarithmic *x*-axis, which visually compresses a wide input range into a manageable space. We show below that the axis choice does not influence results.

The thick dashed lines in Fig. 1B represent the standard deviation of *p*(*r* | *s*), given by *σ*, which is the square root of the response variance. This variance can be expressed as a function of either the signal or mean response,

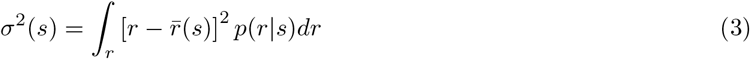

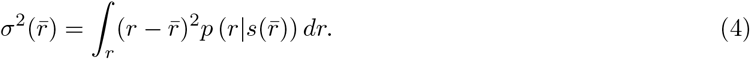

In the latter equation, 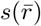 represents the inverse of the dose-response function, meaning that it is the signal value that corresponds to mean response 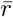. Combining the conditional response distribution with the signal distribution yields the response distribution, *p*(*r*), which represents the probability of receiving a particular response value for an unknown signal value,

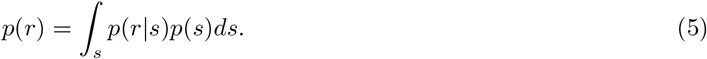

The central focus of this work is on the mutual information between signal and response [48, 56]. In Fig. 1C, the red and blue regions represent the information of the signal and response where, conceptually, each unit of area represents a separate signal or response value. Specific response values that correspond to specific signal values, illustrated by the overlapping region, represent transmitted information and are called the mutual information; this is the amount of information in the response that comes from the signal. There are also signal values that produce identical responses as other signal values. These redundant signals, shown with the non-overlapping red region, do not transmit information. Likewise, there is also information in the response that does not come from the signal but from the noise, shown with the non-overlapping blue region; this information is also not of interest. The mutual information represents the fidelity of the signaling system by quantifying the amount that a cell can learn about the signal value from any individual response value. Its value is [48, 56]

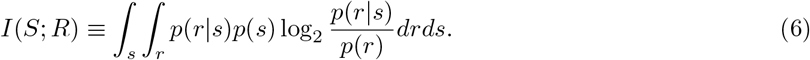

The fact that the logarithm is taken with base 2 implies that the information is quantified in bits, with the interpretation that a cell can reliably distinguish between 2^*I*(*S*;*R*)^ different signal values. For example, 3 bits of information corresponds to 2^3^ = 8 distinguishable signal values.

To compute the mutual information from Eq. 6 requires the signal distribution, *p*(*s*), but it is rarely known for cells in their natural environments [71]. We address this by following the convention of choosing the signal distribution, now written as *p*^*\**^(*s*), as the one that maximizes the mutual information. This optimized mutual information is called the channel capacity and denoted here as *I*^*\**^(*S*; *R*); it is the upper bound to the mutual information. We performed this optimization in several ways. We found it with the small-noise approximation (SNA) for cases where 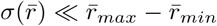, with the large-noise approximation (LNA) for cases where 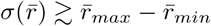, and with the numerical Blahut-Arimoto (B-A) algorithm [56, 69] for all noise values. The approximate analytical methods are described next, and we followed the methods described in [10] for the B-A algorithm.

### Small-noise approximation

Many noise distributions, including Poisson, binomial, and others, can be approximated well by Gaussian distributions if the noise is much less than the mean response range. In this case, *p*(*r*|*s*) is

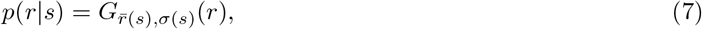

where

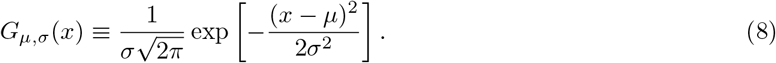

This is a Gaussian with area 1, mean *µ*, and standard deviation *σ*. Tkačik et al. showed that the optimal response distribution for Gaussian noise in the SNA is [69]

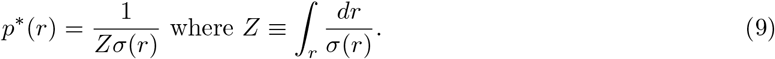

This response distribution can be computed from Eq. 4, using the fact that 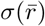 is the same as *σ*(*r*) in the SNA. Tkačik et al. also showed that the channel capacity is [69]

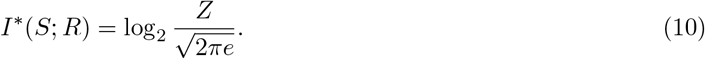

This is exact in the limit of small noise and becomes a lower limit to the channel capacity for larger noise values because the response distribution from Eq. 9 deviates increasingly from the optimal response distribution.

We extended these results to compute the optimal signal distribution. To do so, we used the fact that the dose-response curve relates not just one signal value, *s*, with one mean response, 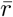, but also the distribution of signals, *p*(*s*), with the distribution of mean responses, 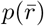; the distribution of mean responses is the response distribution that would arise from the different signal values if there were no noise. This enables a change of variables with the identity (see e.g. section 1.8 of [72]).

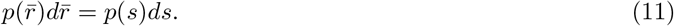

Because of the SNA, the distribution of mean responses equals the response distribution, so 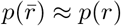. Substituting Eq. 9 into Eq. 11 and simplifying gives the optimal signal distribution,

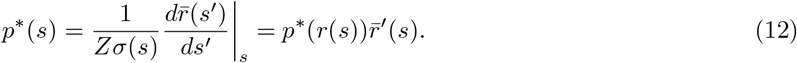

In each equality, the first term increases the likelihood of signals where *σ*(*s*) is small, meaning regions that have low noise, and the second term increases the likelihood in regions where the dose-response curve is steeper, where it better separates different responses and thus improves their discrimination [33].

### Large-noise approximation

No two signals can be discriminated with any certainty if the amount of noise is similar to, or larger than, the range of mean responses. However, the responses do still convey some information about the signals, and this is greatest if the signals, and thus mean responses, are as different as possible [69]. This implies that the optimal signal distribution is a discrete function that restricts the signals to the two extremes, *s*_*min*_ and *s*_*max*_,

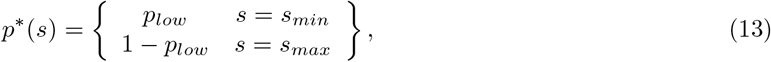

where *p*_*low*_ is the probability of a low signal being sent; its value is given below in Eq. 16. The red graph at the bottom of Fig. 2 illustrates this signal distribution as a pair of delta functions. Each signal results in a separate conditional response distribution, which are shown on the right side of the figure.

**Fig 2.**
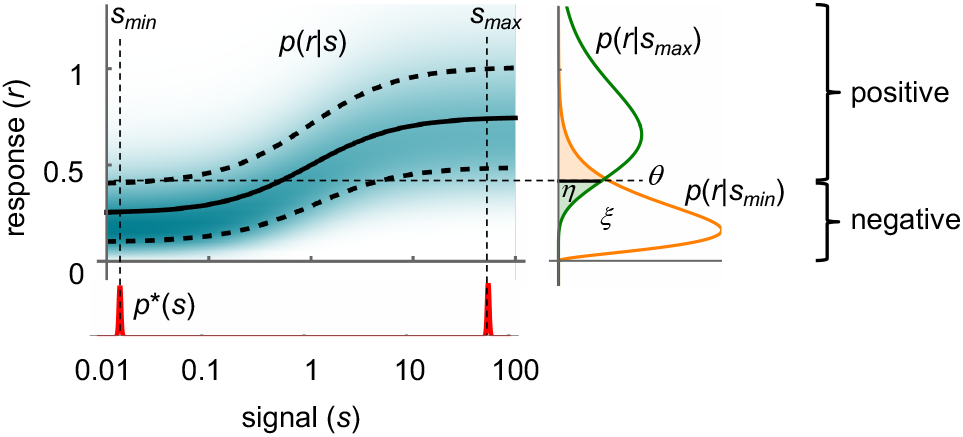
Illustration of large noise approximation method. The signal distribution, *p*^*\**^(*s*), is shown in red at the bottom; it is confined to the signal endpoints, shown near the ends of the signal axis here, but ideally at *s*_*min*_ = 0 and *s*_*max*_ = *∞*. These two signals yield the two conditional response distributions shown on the right. Response values below *θ*, shown with a horizontal dashed line, are recorded as “negative” while those above *θ* are recorded as “positive.” The area of the *p*(*r*|*s*_*min*_) curve that is below *θ*, given as *ξ*, represents the probability of a true-negative and the area of *p*(*r* |*s*_*max*_) that is below *θ*, given as *η*, represents the probability of a false-negative. The mutual information can be computed from *ξ* and *η*. In this illustration, the dose-response function is a Hill function with *A* = 0.5, *B* = 0.25, *E* = 1, and *N* = 1, and the noise is gamma-distributed with scaling factor *n*_Γ_ = 0.3.

Because only two signals are possible, it makes sense to partition the continuous range of response values to two discrete outputs, which we call “negative” if the response is less than some threshold *θ* and “positive” if the response is greater than *θ*. This leaves the question of how to set *θ*. A simple approach is to set *θ* to the response value where the two probability curves cross each other; by symmetry, this is optimal if the probability curves are mirror images of each other, and Tkačik et al. also claimed that it is optimal for unclipped Gaussian probability curves [69] but our work shows otherwise (Appendix). As shown below, the optimal threshold needs to be found numerically in most cases.

Following Tkačik et al. [69], we define *ξ* as the probability of a true-negative, meaning that a low signal was sent and a negative response was observed, and *η* as the probability of observing a false-negative, meaning that a high signal was sent but a negative response was observed. These are shown as areas under probability curves on the right side of Fig. 2, and are given as

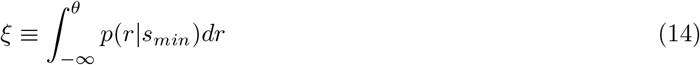

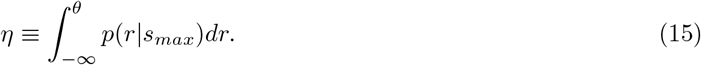

These simplifications reduce the signaling system to the so-called binary asymmetric channel, for which the optimal probability of choosing a low input is [73]

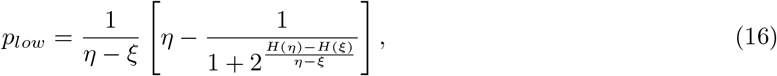

and the channel capacity is

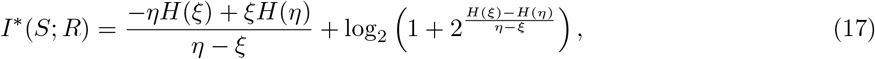

where *H*(*x*) is the entropy function, defined as

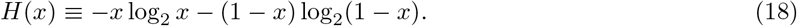

If the probability of a false-negative equals that of a false-positive, meaning that *η* = 1*−ξ*, then the system simplifies to the symmetric binary channel [56] in which

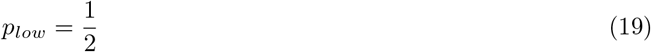

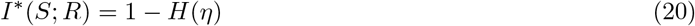

The channel capacities in Eqs. 17 and 20 are necessarily between 0 and 1 bits, and are lower limits to the true channel capacity due to the assumptions of binary signals and responses.

### Summary

For a given signaling system, we assume that one can use experiments or simulations to determine the dose-response function and its variation which, together, represent the conditional response probability, *p*(*r*| *s*). From it, we want to compute the optimal signal distribution, optimal response distribution, and channel capacity. For the small-noise approximation, these results are presented in Eqs. 12, 9, and 10, respectively. For the large-noise approximation, the signal distribution is in Eqs. 13 and either Eq. 16 or Eq. 19, the response distribution can be computed from Eq. 5, and the channel capacity is in either Eq. 17 or 20. If neither approximation holds, then results need to be computed numerically with the Blahut-Arimoto algorithm.

### Signaling over one step

#### Uniform Gaussian noise

In our first computation of the channel capacity for signaling over a single step, we assume that the noise is the same for all signal values (Fig. 3A). This yields analytically solvable results that illustrate the use of the equations presented above. Few biochemical systems, if any, exhibit this type of noise, but it might be a reasonable approximation for some types of laboratory measurements (in this conception, the signaling system starts at a physical cell and ends at its measurement). With the further assumption that the noise is Gaussian with standard deviation *n*_*U*_, the noise standard deviation (Eqs. 3, 4) is

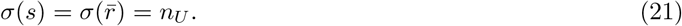

**Fig 3.**
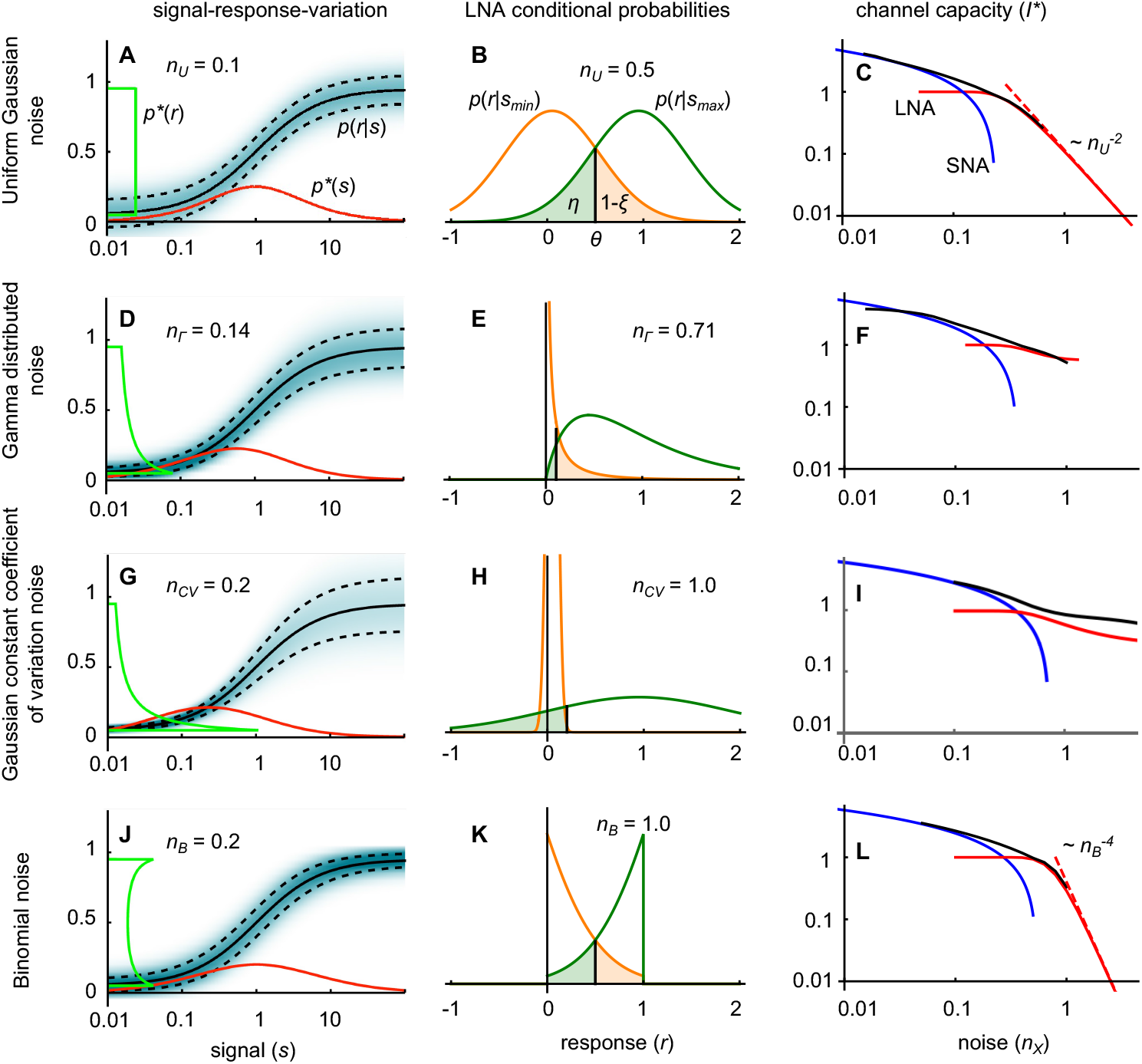
Signaling over one step. SRV plots (left column), conditional response distributions used in the LNA (middle column), and channel capacity computations (right column) for different noise assumptions. Noise assumptions are uniform Gaussian noise (first row), Gamma distributed noise (second row), Gaussian noise with a constant coefficient of variation (third row), and binomial noise (fourth row). SRV plots are depicted as in Fig. 1, and also show the optimal signal and response distributions for the SNA in red and green curves. These plots use a logarithmic signal axis, so *p*^*\**^(*s*) functions are computed by differentiating with respect to ln *s*. In the middle column, conditional response distributions show *p*(*r*|*s*_*min*_) in orange and *p*(*r* |*s*_*max*_) in green, the optimal threshold with a black line, the false negative probability with green shading, and the false positive probability with orange shading. In the right column, channel capacity computations show SNA results in blue, LNA results in red, numerically optimized channel capacities in black, and large-noise trends with red dashed lines (the dotted purple line in panel I, just below the red line, represents LNA calculations with the threshold set at the junction of the two conditional probability distributions). All panels were calculated for 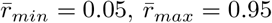. Where necessary, the dose-response curve was set to a Hill function with *A* = 0.9, *B* = 0.05, *E* = 1, and *N* = 1. Noise parameter values are given where appropriate.

Substituting this variation into Eqs. 9, 12, and 10 yields the optimal response distribution, signal distribution, and channel capacity for small noise,

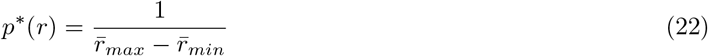

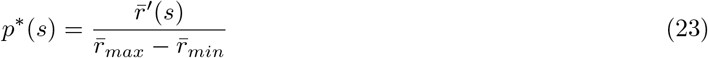

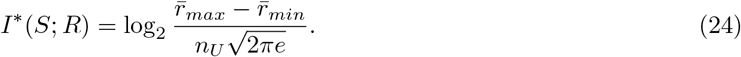

The first equation shows that the optimal response distribution is uniform over the response range (green line in Fig. 3A), which is a well-known result for uniform noise [33, 48]. The second equation shows that the optimal signal distribution is directly proportional to the derivative of the dose-response function, placing the peak of the optimal signal distribution at the steepest part of the dose-response curve (red line in Fig. 3A). The final equation shows that the channel capacity is proportional to the logarithm of the ratio of response range to noise. This makes sense because this ratio is approximately equal to the number of distinguishable signals. Eq. 24 is also similar to the Shannon-Hartley theorem from information theory, which states that the channel capacity scales as the logarithm of the signal-to-noise ratio [56, 74]. A notable aspect of Eq. 24 is that the channel capacity is independent of the shape of the dose-response curve.

With large noise levels, the signals are restricted to *s*_*min*_ = 0 and *s*_*max*_ = *∞*, and the conditional response distributions are restricted to a pair of unclipped Gaussians, each with standard deviation *n*_*U*_ (Fig. 3B). By the symmetry of the system, the optimal probabilities for sending high and low signals are equal (*p*_*low*_ = 1*/*2), the optimal threshold is centered between the mean response values (*θ*^*\**^ = (*r*_*max*_ *− r*_*min*_)*/*2), and the probability of a false negative equals that of a false positive (*η* = 1 *− ξ*). Evaluating the false negative probability using Eq. 15 yields

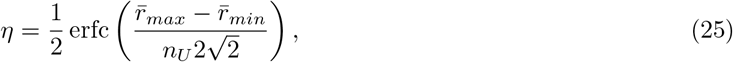

where “erfc” is the complementary error function. Substituting this into Eq. 20 gives the large-noise channel capacity as

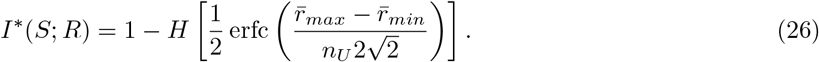

This function has a maximum of 1 bit with small noise, where the LNA is a poor approximation but still a valid lower limit, and decreases as 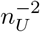 with large noise (red line in Fig. 3C). As for the SNA solution, Eq. 26 shows that the channel capacity depends only on the ratio of response range to noise, and not on the shape of the dose-response curve.

Numerically optimizing the signal distribution, using discretized probability distributions and the B-A algorithm, yielded channel capacity results that agreed well with the analytical equations at low and high noise values and smoothly interpolated between them for intermediate values (black line in Fig. 3C). The optimal signal distribution (Supp. Inf. §1) also smoothly interpolated between the extreme cases, starting with the unimodal distribution of the SNA result, then adding peaks at the extreme values, and finally removing the central peak to create the bimodal distribution of the LNA result.

#### Gamma distributed noise

In equilibrium and many steady-state processes, molecule copy numbers exhibit a Poisson distribution, in which their standard deviations are equal to the square root of their means. Stochastic transcription and degradation of mRNA transcripts is such a process. Amplification of those copy numbers through protein translation maintains the distribution shape but rescales it, producing what it called a gamma distribution [75]. It introduces a scaling factor, *n*_Γ_, to the standard deviation relationship,

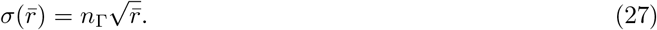

Assuming gamma distributed variation in a one-step signaling system leads to the SRV curve shown in Fig. 3D. With the SNA, the optimal response distribution, signal distribution, and channel capacity are computed by substituting Eq. 27 into Eqs. 9, 12, and 10, leading to

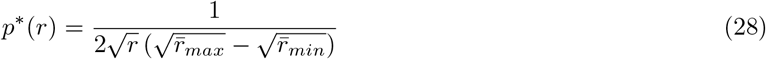

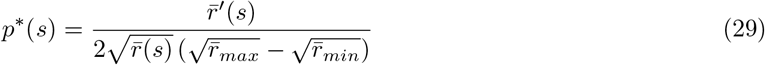

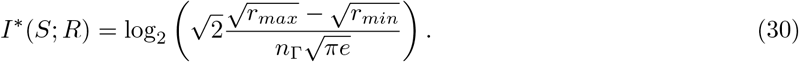

Compared to the uniform noise case, both the optimal response distribution and optimal signal distribution are weighted towards smaller values, due to the lower noise there (Fig. 3D). Likewise, the channel capacity has an increased sensitivity on variation in *r*_*min*_ and decreased sensitivity on *r*_*max*_.

To compute channel capacity values with the LNA, we start with the conditional probability distribution,

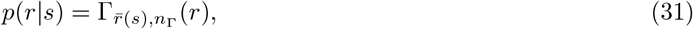

where Γ_*µ,n*_(*x*) is the gamma distribution with mean *µ* and scaling coefficient *n*. Substituting in the extreme signal values *s*_*min*_ = 0 and *s*_*max*_ = *∞* leads to the two conditional response distributions shown in Fig. 3E.

While we were able to analytically integrate these equations to yield the *ξ* and *η* values as functions of *θ* using Eqs. 14 and 15 (Supp. Inf. §2), we were only able to optimize *θ* numerically, which the figure shows with a black line. We then computed the channel capacity with Eq. 17. The two conditional probability distributions are very different from each other, of which *p*(*r* | *s*_*min*_) is highly peaked and *p*(*r*| *s*_*max*_) is quite broad; these differences arise from the dependence of the standard deviation on the response, Eq. 27. As a consequence, the channel capacity decreases slowly with increasing noise (Fig. 3F). This makes sense because, even with high noise, a response that arises within the narrow peak has high probability of arising from a low signal, and a response outside of this region has a low probability of arising from a low signal.

As before, B-A channel capacity calculations smoothly interpolate between the SNA and LNA results (Fig. 3F).

#### Constant coefficient of variation

Highly expressed proteins generally exhibit cell-to-cell variation that depends to a greater extent on the mean expression, and is better modeled as having a constant coefficient of variation [10, 30, 52, 75–77]. The standard deviation of this “constant-CV” noise is directly proportional to the mean response,

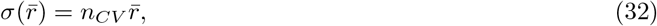

where *n*_*CV*_ is the constant of proportionality (this constant is typically represented as *η*, but we use *n*_*CV*_ here for consistency with the rest of this work).

This constant-CV distribution continues the changes described above for gamma distributed noise. The SRV curve (Fig. 3G) has a greater noise difference between low and high responses, so the SNA optimal signal and response distributions are shifted further toward low signal and response values. Following the same procedure as before, the SNA optimal response distribution, signal distribution, and channel capacity are

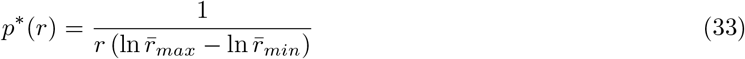

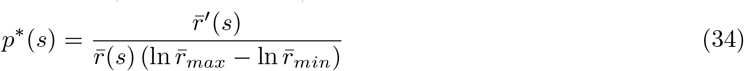

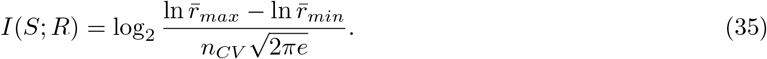

This channel capacity is more sensitive to the response baseline, formally approaching infinite information transmission as the baseline of the dose-response curve decreases toward zero (based on the unrealistic assumption that the noise decreases to zero as well).

As an aside, constant-CV distributions are convenient to use because the logarithmic terms in Eqs. 33 to 35 simplify so that only the ratio of 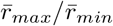 is required. As a result, one doesn’t need to know actual response molecule counts or other absolute values [10].

For large noise, we assume that the conditional probabilities are Gaussian distributed and we allow negative responses. These distributions integrate to yield *ξ* and *η* (Supp. Inf. §3), but require numerical optimization to position *θ*. As with the gamma distribution, the extreme conditional probability distributions have one narrow peak and one broad peak (Fig. 3H), causing the channel capacity to decrease slowly with increasing noise (Fig. 3I). However, the fact that we allow negative responses reveals a shortcoming of the LNA approach. Because *p*(*r* | *s*_*max*_) extends to both sides of *p*(*r* | *s*_*min*_), one would ideally define a threshold on each side of the narrow peak so that only responses inside both thresholds would be considered “negative”. We only use one threshold, which explains why Fig. 3I shows that the LNA channel capacity does not approach the B-A channel capacity for high noise levels. Nevertheless, as always, the LNA result is a valid lower limit.

#### Binomial noise

Binomial noise arises when information is stored in the state of a two-state chemical species, such as ligand-binding by receptors, protein phosphorylation at a single site, opening or closing of ion channels, and activation or repression of transcription.

Labeling the two states as “active” and “inactive”, if a protein has probability *P* of being active, then the probability of finding *k* active proteins, out of *M* total proteins, is given by the binomial distribution and has response standard deviation of 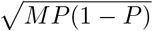 [78]. We scale this for a system in which a response value of 1 corresponds to full activity. In this case, 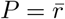 and the standard deviation is

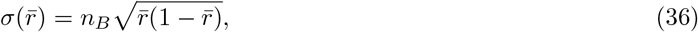

where *n*_*B*_ = *M* ^*−*1*/*2^. This use of a noise coefficient, *n*_*B*_, maintains consistency with the notation used above and also allows the coefficient to be varied to account for downstream noise amplification.

The SRV curve for binomial noise (Fig. 3J) shows that the noise is small for low and high response values and large for intermediate responses. As a result, the optimal response and signal distributions, also shown in the figure, spread out toward the ends of their ranges. Computing the SNA optimal response distribution, signal distribution and channel capacity yields

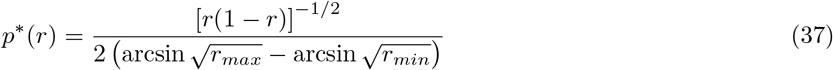

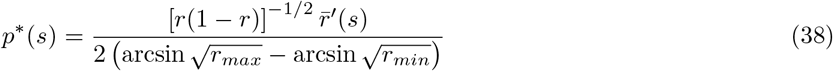

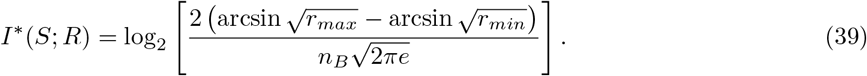

These equations simplify substantially if there is no basal activation and no attenuation of maximal activation, meaning that *r*_*min*_ = 0 and *r*_*max*_ = 1,

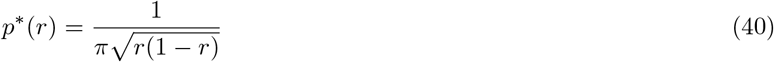

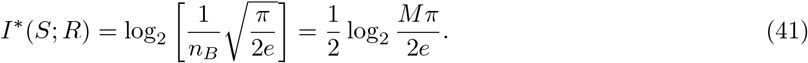

The latter equation shows that the channel capacity scales as the logarithm of the number of two-state signaling proteins. For example, 30 proteins are sufficient for 2 bits of information transmission, 60 proteins for 2.5 bits, and 120 proteins for 3 bits. This leads to the prediction that cells are likely to express far fewer proteins to detect binary signals than continuous ones [79]. This can also be seen as the minimum metabolic cost to achieve a given level of signaling fidelity when using two-state proteins.

To compute large noise results, which apply when there are few total two-state proteins or when binomial noise gets amplified, we defined a continuous version of the binomial distribution (Supp. Inf. §4). It simplifies the computation of the conditional response distributions with the extreme inputs, *p*(*r*|*s*_*min*_) and *p*(*r* | *s*_*max*_), shown in Fig. 3K. These distributions have a high amount of overlap, and this overlap only increases with increasing noise, which creates a rapid decline in channel capacity values with increasing noise levels (Fig. 3L). We empirically found that the LNA channel capacity scales as 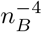. As usual, B-A channel capacity calculations interpolated smoothly between the SNA and LNA results.

#### One-step signaling example

We illustrate the use of these equations by estimating how much information is transmitted when a stimulus is added to a population of cells where the “noise” is the cell-to-cell variation in responses. Starting with the noise term, the variation between copy numbers of proteins that are expressed at low or intermediate rates have been shown to follow gamma distributed statistics [75, 76, 80]. Bar-Even et al. quantified the noise parameter in yeast cells as 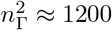 [76] (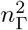 is in units of copy numbers here, whereas a scaled version was used in Fig. 3D-F). Supposing that a stimulus causes a response protein to increase from about 1200 copies to about 12,000 copies, where the latter number is a rough average of steady-state protein amounts from Fig. 2 of the same paper [76] and the former number assumes 10% basal response, we can estimate the channel capacity using the results presented above for the gamma distribution. The SNA result (Eq. 30) is about 0.6 bits, the LNA result is about 0.9 bits, and the B-A result is 1.1 bits, which is the most accurate answer. This final value agrees well with the roughly 1 bit that has been found in multiple experiments [10, 24, 52, 81, 82].

#### One-step signaling dose-response curves

For all four noise distributions, the SNA channel capacity equations depend only on *r*_*min*_, *r*_*max*_, and the noise parameter, and not on the shape of the dose-response curve. This latter independence turns out to be a general result, including for all possible noise distributions and amounts, which can be seen by discretizing the mutual information integral (Eq. 6) to the sum

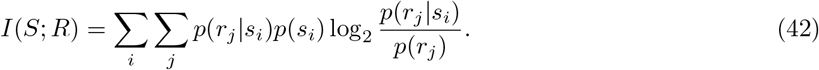

The *s*_*i*_ and *r*_*j*_ vectors replace the *s* and *r* variables and the *p*(*s*_*i*_), *p*(*r*_*j*_), and *p*(*r*_*j*_ |*s*_*i*_) vectors and matrices replace the *p*(*s*), *p*(*r*), and *p*(*r* |*s*) functions. This equation, which is typically given as the definition of mutual information for discrete systems [48, 56], is independent of the signal and response values, along with the shape of the dose-response curve, but depends only on their probability distributions. As a practical matter, this independence of the mutual information on specific signal values also means that the choice to represent signals with either linear or logarithmic axes does not affect channel capacity results. Likewise, discretizing the response distribution integral, Eq. 5, shows that optimal response distributions are also insensitive to dose-response curve shapes.

On the other hand, for low or moderate noise, the optimal signal distribution does depend on the dose-response curve shape (see Eq. 12). Its peak is generally near the steepest part of the dose-response curve and is also biased toward responses that exhibit less noise. For a Hill function dose-response curve, we found that the optimal signal distribution’s peak is at the EC_50_ in both the uniform Gaussian noise case and the binomial noise case, and at about half of the EC_50_ for gamma distributed noise (Supp. Inf. §5). The peak is further shifted to lower concentrations, and is highly sensitive on the *r*_*min*_ value, for constant-CV distributed noise.

However, biological cells cannot necessarily optimize their signal distributions. For example, humans and other animals both secrete and sense the epinephrine signaling molecule when under stress, but the frequency of being under stress is not something that we optimize, and certainly not for the purpose of achieving optimal signaling. If one assumes instead that there is some given signal distribution, then one would expect that cells would adapt their dose-response curves to match the natural signal distribution [33]. The optimal dose-response curve can be found in the SNA by inverting Eq. 12 to yield

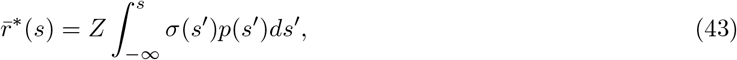

where *Z* can be computed from Eq. 9 or just treated as a normalization parameter.

With high noise, the optimal signal distribution is limited to just the minimum and maximum input values (Eq. 13), making it insensitive to the dose-response curve shape.

### Signaling over two steps

#### Two step system

To investigate information transfer over two steps, we add an intermediate node, “A”, to the signaling system, so now a signal goes from S to A in step 1, and from A to R in step 2 (Fig. 4A). Information traverses the pathway sequentially, so the mutual information through the whole pathway, from S to R, can’t exceed that for either step individually (Fig. 4B). This is called the data processing inequality in information theory [56] and simply reflects the fact that information from S to R has to go via A.

**Fig 4.**
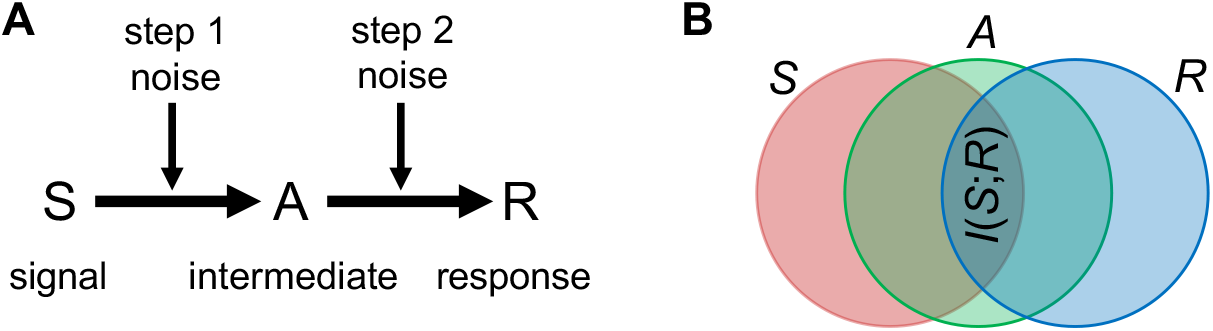
Signaling over two steps. (A) Diagram of two-step signaling. (B) Venn diagram showing information loss over the pathway. The red region represents signal information, the green region is information at A, and the blue region is response information. Overlaping regions represent mutual information.

How can a cell optimize its signaling system in order to transmit the most information? Starting with step 1, the results presented above show that cells transmit more information with lower response baselines, wider response dynamic ranges, and dose-response curves that are tuned to the natural signal distribution. These also apply to step 2, but now with the constraint that the step 1 response distribution is the step 2 signal distribution. This distribution is already fully optimized through the step 1 response range and noise distribution, so the only option is that the step 2 dose-response curve needs to match the given signal distribution. We argued previously that this second dose-response curve, or the transfer function, should be linear in order to optimize information transmission [43, 45–47]. Here, we confirm those results for some situations and extend them for others.

Fig. 5 presents a qualitative illustration of why a linear transfer function improves information transmission for systems that have relatively low noise. Panel A shows step 1, in which 5 different input signals, each of which starts with a well-defined value, get converted to 5 intermediate responses that are less well defined; each is the conditional probability value, *p*(*a* | *s*), for the given signal. The input signal values were chosen in this example so that the intermediate responses are well spaced out, roughly in accordance with the optimal response distribution. These intermediate responses become the input to step 2, which is shown with a linear transfer function in Panel B and a nonlinear transfer function in Panel C. In step 2, the signals gain more noise, leading to broader output distributions. More importantly, they stay well spaced out if the transfer function is linear, which keeps them distinguishable from each other, but multiple ones overlap each other if the transfer function is nonlinear; this reduces distinguishability and indicates loss of mutual information.

**Fig 5.**
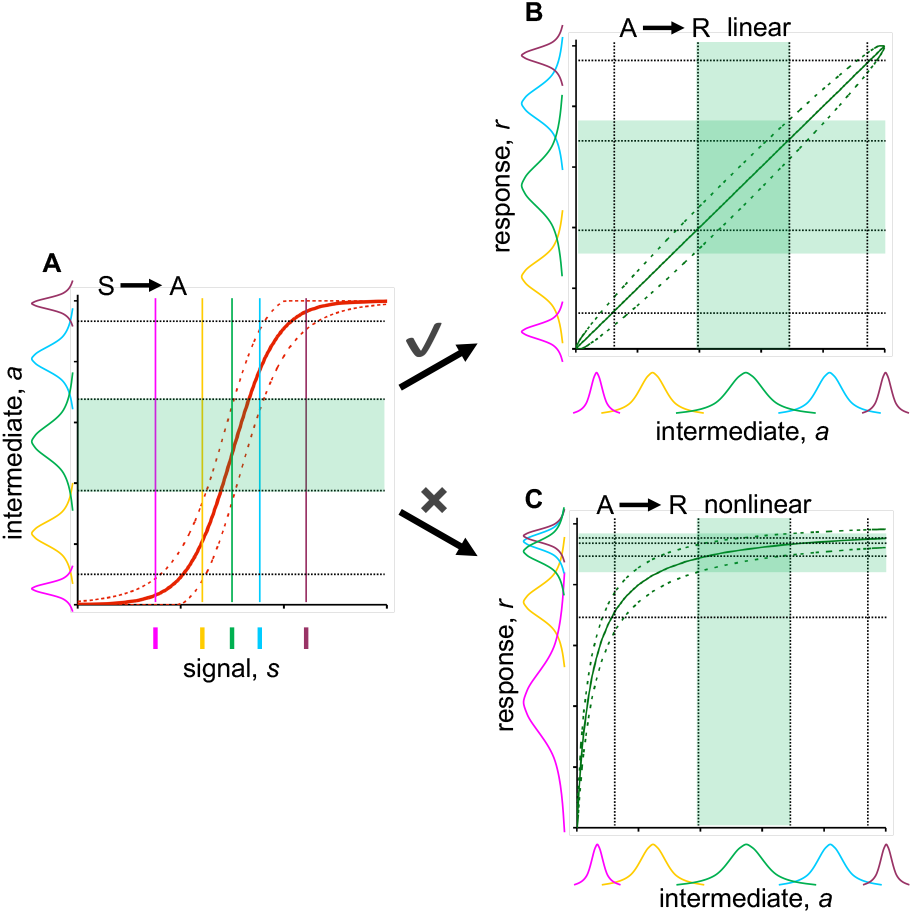
Signaling over two steps with linear and non-linear transfer functions. (A) Illustration of signaling over the first step, in which 5 different color-coded input signals yield 5 noisy but still distinguishable intermediate outputs; shading in the figure represents the noise introduced for the central green input signal. (B) Illustration of signaling over the second step, assuming a linear transfer function. Each of the original input signals gains additional noise here, but the final output response distributions are still uniformly spaced and so are mostly distinguishable. (C) Another illustration of signaling over the second step, this time with a non-linear transfer function. Now, several of the output response distributions overlap each other, which makes these signals indistinguishable and represents lower information transmission. While it doesn’t affect results or interpretation, note that the panel A *x*-axis is logarithmic and all other axes are linear.

#### Small noise approximation

To investigate 2-step signaling more rigorously, we start by assuming that noise is small. The 2-step system is defined by the conditional probabilities for the separate steps, *p*(*a* |*s*) and *p*(*r* |*a*), along with the signal distribution *p*(*s*). These yield the dose-response functions for the two separate steps, 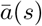 and 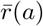, using Eq. 1 with suitable variable substitutions. Due to the SNA, we are allowed to simplify our notation by dropping the overbars in these functions, which we do throughout the rest of this section.

The pathway dose-response function is the composition of the single-step dose-response functions,

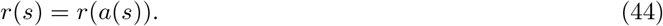

We illustrate this with two examples. First, composing an arbitrary step 1 dose-response function, *a*(*s*), with the linear transfer function *r*(*a*) = *c*_1_ + *c*_2_*a*, where *c*_1_ and *c*_2_ are arbitrary constants (Fig. 5B, leads to

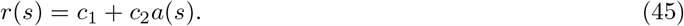

This only differs from *a*(*s*) in that it introduces a baseline and performs rescaling. For example, if *a*(*s*) is a Hill function, then *r*(*s*) is another Hill function with the same EC_50_ and Hill cooperativity, showing dose-response alignment. Second, if both *a*(*s*) and *r*(*a*) are Hill functions (Fig. 5C), it turns out that EC_50_ of the composition is necessarily smaller than that for *a*(*s*). This means that the dose-response curve for the pathway is left-shifted compared to that for the first step [46, 83] and there is no dose-response alignment.

We compute the noise term for the entire pathway, *σ*(*r*), by first considering only the noise that arises in step 1. This step 1 noise is *σ*_1_(*a*) at node A, where we’re expressing it as a function of the downstream node value (see Eq. 4). Transmitting this noise to R transforms it due to the slope of the *r*(*a*) dose-response curve (see Fig. 5C), causing the step 1 noise at R to be

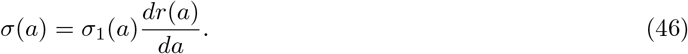

Rewriting this as a function of the response value yields

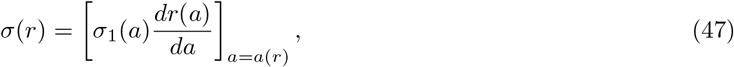

where *a*(*r*) is the inverse of the dose-response function *r*(*a*). The derivative of a function is equal to the reciprocal of the derivative of the inverse function (i.e. *dy/dx* = (*dx/dy*)^*−*1^), enabling the rearrangement

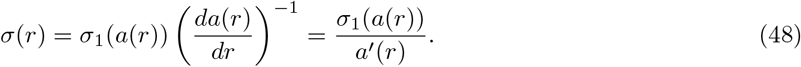

In the second equality, the prime symbol indicates a derivative with respect to *r*. In addition to this noise from step 1, there is also noise that arises in step 2, which is given as *σ*_2_(*r*). Assuming that these noise sources are independent from each other, then the sum of their variances is the total variance at R,

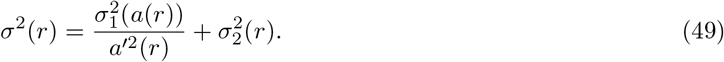

Next, we optimize the channel capacity for signaling over two steps using this total noise distribution. Because we assume the SNA, Eq. 10 gives the channel capacity for signaling from S to R, where *Z* is defined in Eq. 9. The channel capacity increases monotonically with *Z*, so we save some steps by searching for the dose-response function that maximizes *Z*, knowing that it also maximizes *I*^*\**^(*S*; *R*). Expanding *Z* in terms of the pathway noise (Eq. 49) yields

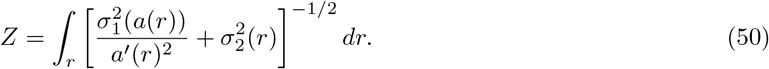

Using the method of the calculus of variations, an approach that finds the function that produces an extremal value (see e.g. [84, 85]), we define *F* (*a, a’*, *r*) as the integrand of *Z*,

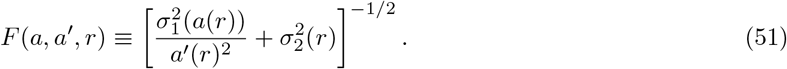

The Euler-Lagrange differential equation can now be solved to identify the conditions that maximize *Z*. It is

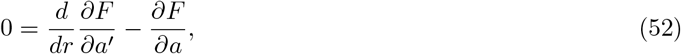

where

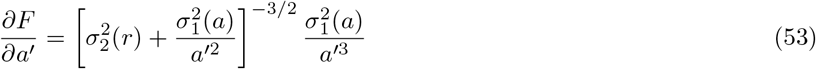

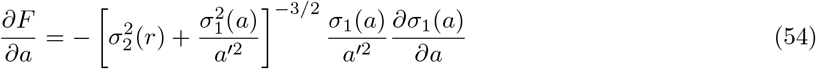

(we dropped the explicit dependence of *a’* and *a* on *r*).

We need a noise distribution at this point, so we assume uniform Gaussian noise for both steps, meaning that *σ*_1_(*a*) = *n*_*U*1_ and *σ*_2_(*r*) = *n*_*U*2_ (see Eq. 21). Substituting these into the Euler-Lagrange differential equation, Eq. 52, yields

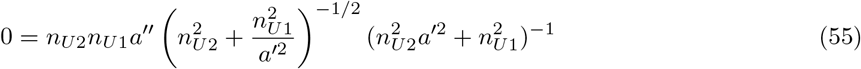

Inspection shows that the right hand side can only equal zero if *a*″ = 0. This is a trivial differential equation, which solves to

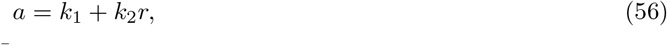

where *k*_1_ and *k*_2_ are constants of integration. Inverting gives

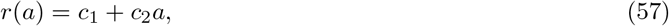

where *c*_1_ and *c*_2_ are now some other constants (equal to *−k*_1_*/k*_2_ and 1*/k*_2_, respectively). Thus, we have proven, in the case of uniform noise, that the channel capacity is maximum when the transfer function is linear, with an optional baseline offset (*c*_1_) and scaling factor (*c*_2_).

We can now use this solution to solve for the total noise in the pathway and then the channel capacity. The total noise is given by Eq. 49, into which we substitute the solution for *a’* (*r*) from Eq. 56, while exchanging *k*_2_ for 1*/c*_2_, and we also substitute in the single-step noise terms from Eq. 21 for *σ*_1_(*a*) and *σ*_2_(*r*) to yield

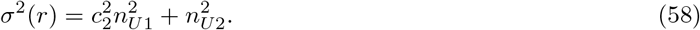

This is a constant, so the entire pathway still exhibits uniform Gaussian noise, with this variance (it’s Gaussian because both terms arise from Gaussian distributions). Finally, this noise is substituted into Eq. 24 to yield the channel capacity for the complete 2-step pathway as

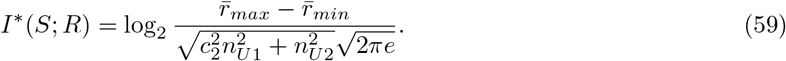

Note that the total pathway noise in Eq. 58 is necessarily larger than the step 2 noise by itself and, correspondingly, the pathway channel capacity is less than the step 2 channel capacity, both of which make sense and agree with the data processing inequality. It initially appears that one could completely remove the effects of the step 1 noise by making *c*_2_ small enough. Indeed, this is true, but it’s not that simple; *c*_2_ is the slope of the transfer function, so it equals

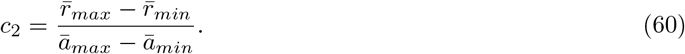

Reducing *c*_2_ through the numerator would require a smaller response dynamic range, which would have the net effect of reducing channel capacity due to its impact on step 2 (i.e. it is the numerator of Eq. 59). On the other hand, reducing *c*_2_ through the denominator would require a larger intermediate species dynamic range, which might be achievable, but this is just the same dynamic range effect that we saw before in Eq. 24.

The same procedure works for the other noise distributions. As a second example, suppose both step 1 and step 2 use the constant-CV distribution. In this case, the Euler-Lagrange differential equation (Eq. 52) solves to give

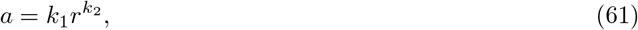

which inverts to give

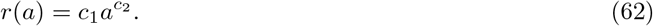

A linear transfer function is still an optimal solution, but now the transfer function cannot have a baseline offset. Instead, it can be proportional to a power of its input so, for example, *r* could be proportional to *a*^1*/*2^ or to *a*^2^ without any reduction in channel capacity. As in the prior example, we can use this solution to find the total pathway noise, which is

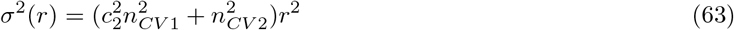

and this can then be substituted into Eq. 35 to yield the channel capacity for the pathway as

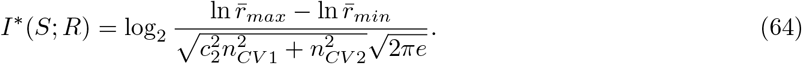

Table 1 lists optimal transfer function shapes for the uniform, gamma, and constant-CV noise distributions in each of the two steps (see Supp. Inf. §6 for binomial noise distributions). The table shows that if the two steps have the same noise distribution, then direct proportionality is always an optimal solution. However, due to the two constants of integration, it is only one of a family of equally optimal transfer functions. On the other hand, direct proportionality is never an optimal solution if the two steps have different noise distributions.

**Table 1.** Optimal transfer functions for low noise signaling.

| step 1 \ step 2 | Uniform | Gamma | Const. CV |
| --- | --- | --- | --- |
| Uniform | $c_1 + c_2 a$ | $c_1 (c_2 + a)^2$ | $c_1 e^{c_2 a}$ |
| Gamma | $c_1 + c_2 \sqrt{a}$ | $c_1 (c_2 + \sqrt{a})^2$ | $c_1 e^{c_2 \sqrt{a}}$ |
| Const. CV | $c_1 + c_2 \ln a$ | $c_1 (c_2 + \ln a)^2$ | $c_1 a^{c_2}$ |

These are the optimal transfer functions for 2-step signaling in the SNA; they are also given as *r*(*a*) and described as the step 2 dose-response functions. The *c*_1_ and *c*_2_ values are arbitrary constants of integration. Note the diagonal elements can always simplify to *r*(*a*) = *a* with suitable choices of *c*_1_ and *c*_2_.

### Large noise approximation

If there are high levels of noise, then the optimal signal distribution changes to a pair of delta functions at the two extreme signal values, *s*_*min*_ and *s*_*max*_ (Eq. 13 and bottom of Fig. 6A). This creates intermediate responses, *p*(*a* |*s*_*min*_) and *p*(*a* |*s*_*max*_), that are as separated as possible, and hence maximally distinguishable from each other (left side of Fig. 6A). These response distributions overlap some, which can be quantified with the true and false negative probabilities, *ξ*_*a*_ and *η*_*a*_ (Eqs. 14 and 15). They are also the signal distributions for step 2, where they accumulate additional noise to become the final response distributions, *p*(*r*|*s*_*min*_) and *p*(*r*|*s*_*max*_), shown on the left side of Figure 6B.

**Fig 6.**
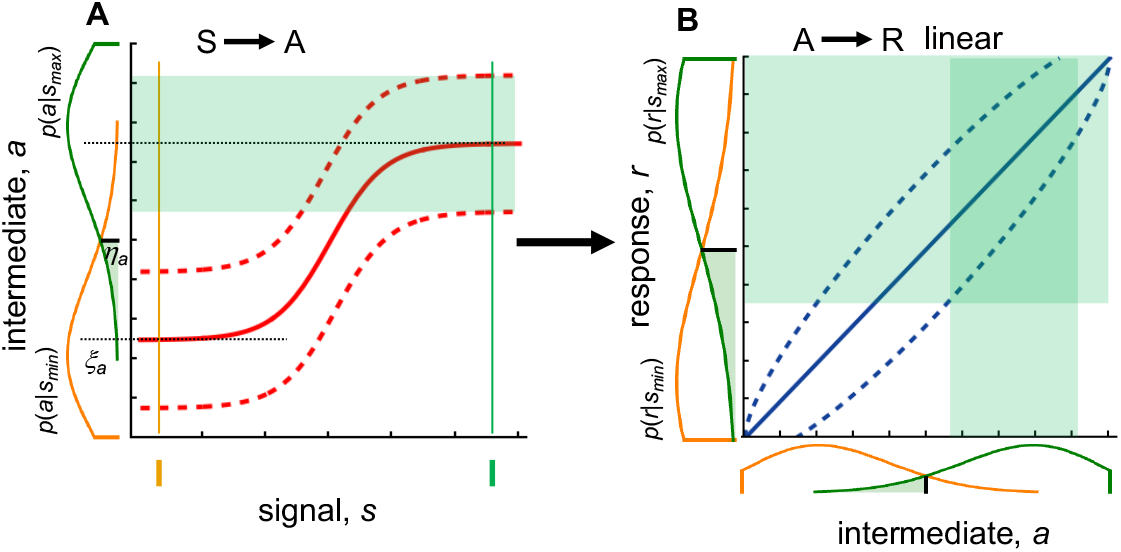
Signaling over two steps with high noise. (A) Illustration of signaling over the first step, in which the two color-coded input signals yield two noisy and partially distinguishable intermediate outputs; shading in the figure represents the noise introduced for the right-hand green input signal. (B) Illustration of signaling over the second step, assuming a linear transfer function. Each of the original input signals gains additional noise here. The panel A *x*-axis is logarithmic and all other axes are linear.

It is straightforward to give equations for the final distributions,

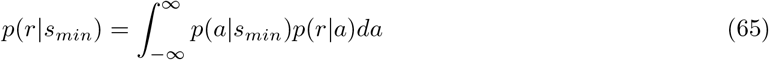

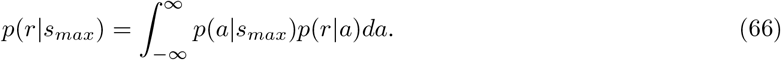

Substituting these into Eqs. 14 and 15 yield the overall true and false negative probabilities, *ξ* and *η*, and then Eq. 17 gives the channel capacity. However, further simplification is not possible without making assumptions about the transfer curve shape and the noise distribution and, even then, results are difficult to interpret. We did not pursue analytical approaches further.

The qualitative reasoning described above in reference to Figure 5 applies equally well to large noise amounts as small noise amounts. It still supports the notion that linear transfer functions typically keep the conditional response distributions as separated as possible and thus transmit more information than non-linear transfer functions. Our simulations, described below, also support this conclusion.

### Switchlike transfer functions

Switchlike transfer functions, which are an extreme version of ultrasensitivity [86–90], represent a very different type of dose-response curve. Here, any input below some switching threshold produces a “low” output and any higher input produces a “high” output. The fact that there are only two outputs means that a maximum of one bit of information can be transmitted.

Starting with the input signal, the optimal signal distribution clearly needs to include signals above and below the threshold (bottom of Fig. 7A). There is no benefit to including more than these two inputs because any additional ones represent redundant information. These two inputs create two intermediate responses that include some noise. The fact that there are only two inputs means that we only need to distinguish between their two responses, which is relatively easy, even with moderate noise. In other words, it’s possible for the same false positive and false negative probabilities that we used before, 1 *− ξ* and *η* (Eqs. 14 and 15), to be essentially zero even when signaling is moderately noisy.

**Fig 7.**
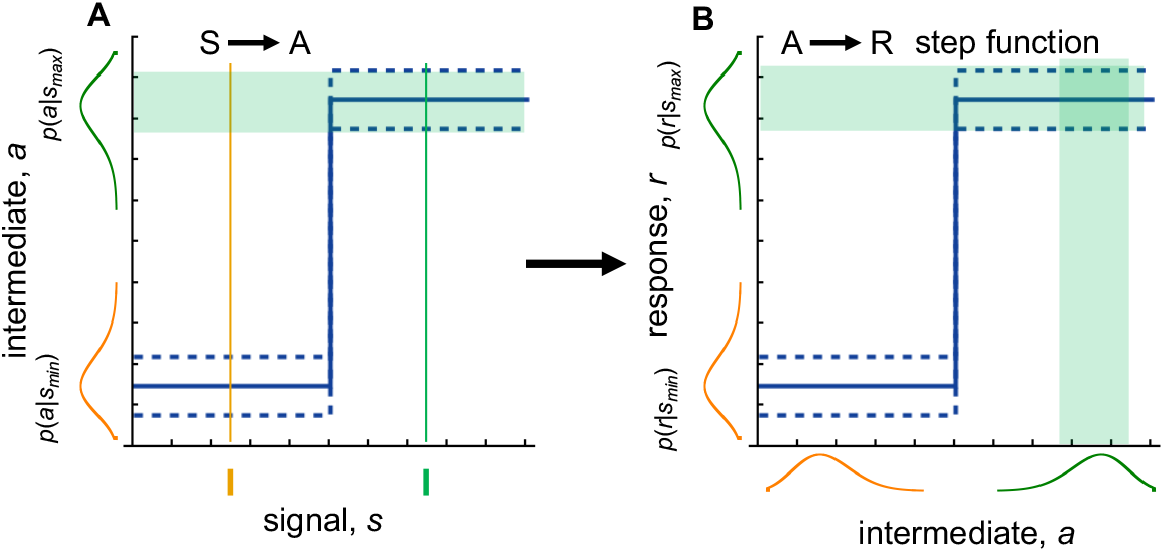
Signaling over two steps with switchlike dose-response functions. (A) Illustration of signaling over the first step, in which the two color-coded input signals yield two noisy and intermediate outputs; shading in the figure represents the noise introduced for the right-hand green input signal. (B) Illustration of signaling over the second step, assuming a switchlike transfer function.

The intermediate responses become the inputs to step 2 (bottom of Fig. 7B). So long as the step 1 noise is small enough that the intermediate distributions don’t cross the switching threshold, then this noise is not propagated downstream due to the fact that the dose-response curve is flat throughout both input regions. As a result, the final response noise is solely the noise that arises from step 2, with no noise from step 1. This process can be repeated indefinitely in a chain of switchlike transfer functions, each one of which removes the prior noise and introduces only its own noise. In this manner, switchlike signaling can transmit 1 bit of information indefinitely without information loss.

On the other hand, if the intermediate distributions do cross the switching threshold, then there is some finite probability of an incorrect response. In this case, the input noise does not create a proportional increase of response noise, as it does with the graded dose-response curves that we investigated above (see Figs. 5 and 6 and Eq. 46), but a response that is completely incorrect. In other words, switchlike signaling is very robust to noise up until the signal distributions cross the switching threshold, but then loses information rapidly as the noise increases beyond that.

To quantify information loss in a two-step system with switchlike dose-response curves, we start with the step 1 true and false negative probabilities, *ξ*_1_ and *η*_1_ (Eqs. 14 and 15). From them, the conditional response distributions are

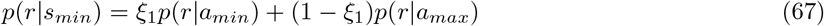

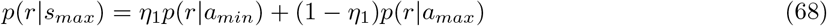

Substituting these results back into Eqs. 14 and 15 leads to the overall true and false negative probabilities,

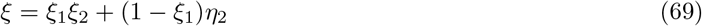

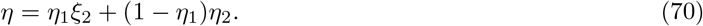

Here, *ξ*_2_ and *η*_2_ are the true and false negative probabilities for the second step alone (quantified by assuming a simple input of two well-defined signals, as shown at the bottom of Fig. 7A). Finally, the channel capacity can be computed from Eq. 17.

As before, these results simplify substantially in the special case that the probability of a false negative equals that of a false positive for both steps (i.e. *η*_1_ = 1 *− ξ*_1_ and *η*_2_ = 1 *−ξ*_2_). This symmetry gets passed on to the full pathway, where

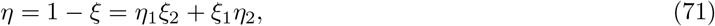

and then the channel capacity is simply 1 *− H*(*η*), from Eq. 20. As an example, consider a symmetric system with a 2% false negative and false positive probability at each step (i.e. *η*_1_ = *η*_2_ = … = 0.02 and *ξ*_1_ = *ξ*_2_ = … = 0.98). The channel capacity drops off rapidly, with 0.86 bits after the first step, 0.76 bits after the second step, and 0.68 bits after the third step.

### Multistep signaling simulations

We investigated information transmission for three toy signaling systems (left column of Fig. 8). In each system, the signal, S, catalytically activates node A, which activates node B, which activates node R. System I uses uncatalyzed inactivation, with the consequence that each node’s dose-response curve is left-shifted and has a lower amplitude than that of the prior node [46, 83]; these imply a non-linear transfer function. In System II, the “inactive” form of each node inactivates the downstream node in a “push-pull” mechanism, creating perfect alignment of the dose-response curves [46]; here, the transfer function is linear. System III adds positive feedback within each node to create switchlike signaling [91]. We ran stochastic simulations of each network using either a low-noise approach, where each node was composed of 100 molecules, or a high-noise approach, where each node had 10 molecules and also had an uncatalyzed reaction that increased the node’s basal activation level (see Methods). After each simulation reached steady-state, we recorded the numbers of active A, B, and R molecules repeatedly. By the ergodic hypothesis, the distribution of these active molecules over time is equivalent to their distributions over multiple simulations.

**Fig 8.**
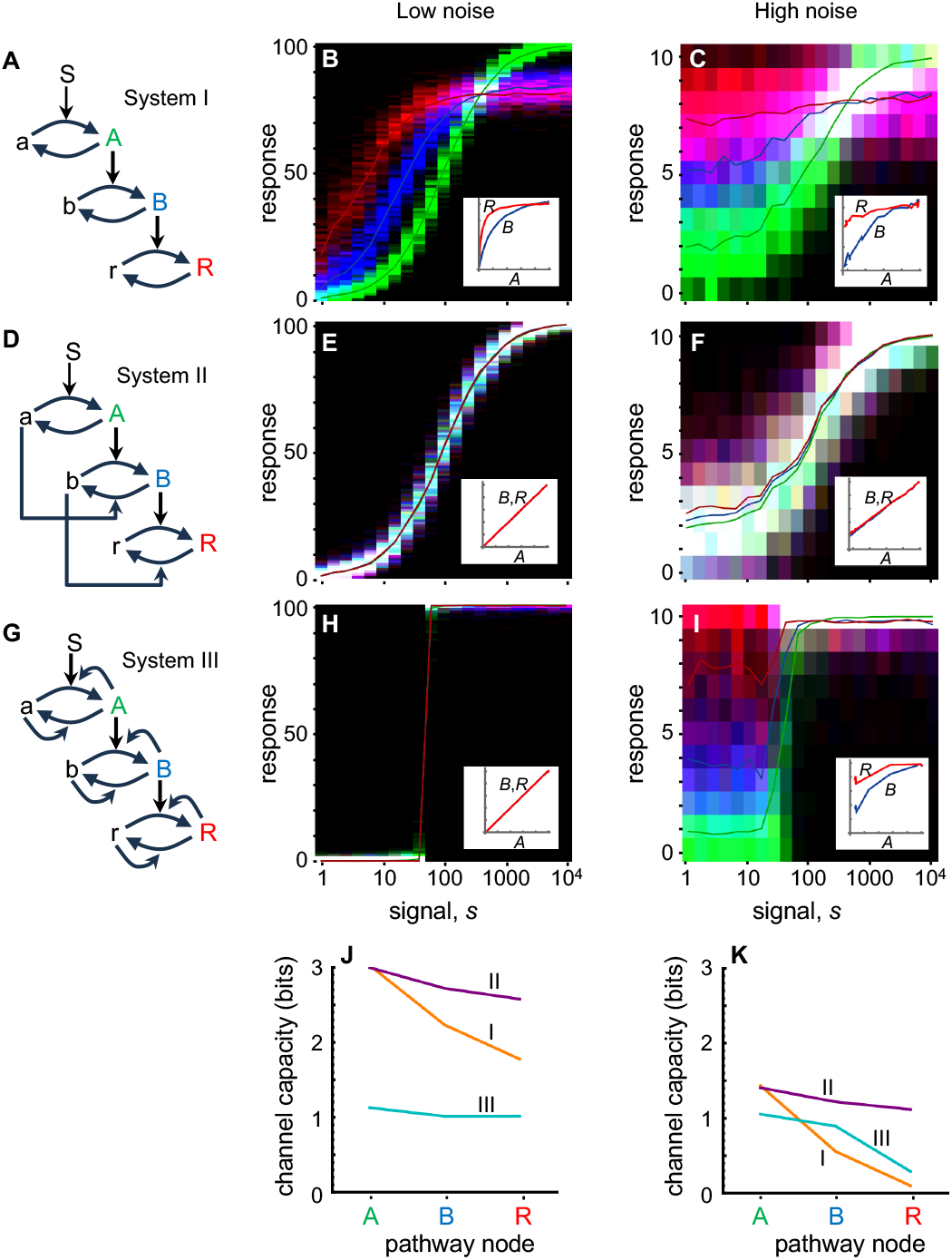
Multistep signaling simulations. (A,D,G) Diagrams of Systems I, II, and III; System I only includes forward activation, System II adds push-pull control, and System III adds stabilizing positive feedback. (B,E,H) “Low-noise” simulation results for the respective systems. Each vertical slice represents a separate simulation, each with the specified number of S molecules. Green, blue, and red regions represent numbers of A, B, and R molecules, respectively, over many time points that are all at steady-state; solid lines depict their mean values. Insets show the transfer functions from A to B in blue, and A to R in red. (C,F,I) “High-noise” simulation results for the same systems, shown in the same manner. (J,K) Channel capacity values at each node, computed with the B-A algorithm, for the low-noise and high-noise simulations, respectively. System I is in orange, System II in purple, and System III in cyan. See Methods for details.

The middle and right columns of Fig. 8 show the simulation results. In each figure panel, each vertical slice represents a single simulation with the given signal amount; the green, blue, and red colors represent the numbers of A, B, and R molecules at each sampling point. These figures, which are essentially SRV plots, also show the time-averaged mean numbers of A, B, and R molecules with solid lines.

The low-noise results (middle column) show the expected results that the System I dose-response curves shift to the left and to lower amplitude for downstream nodes, the System II dose-response curves are perfectly aligned, and the System III dose-response curves are switchlike. The high-noise results (right column) show substantially larger variation for each system and also progressively greater baseline activity for downstream steps in the pathways due to the uncatalyzed activation reactions.

We used the B-A algorithm to numerically compute the channel capacity from the input signal to nodes A, B, and R for each SRV curve (Fig. 8 bottom panels) and found results that agreed well with predictions. For Systems I and II, the channel capacity to node A, where the systems are still identical, is 3.00 bits in the low-noise case and 1.43 bits in the high noise case; the former value agrees with the SNA predicted value of 2.93 bits from Eq. 41 and the latter is consistent with the SNA prediction of 0.81 bits from Eq. 39 and the LNA prediction of 1.00 bits from Eqs. 14 to 17, both of which are always lower limits and known to be inaccurate for situations with intermediate noise levels. Moving downstream, the channel capacity values decrease for nodes B and R, as expected from the data processing inequality. For the low-noise simulations of System II, the channel capacities at nodes B and R are 2.72 and 2.58 bits. These are in reasonable agreement with the SNA predictions for two-step signaling using the binomial noise equivalents of Eqs. 59 and 64 (presented in Supp. Inf §6), which predict 2.43 and 2.13 bits. Meanwhile, System I loses substantially more information than System II does, which agrees with our finding that linear transfer functions are better at preserving information. The high-noise simulations show the same trends, with System II losing a small amount of information as the signal moves downstream and System I losing a large amount of information.

The System III channel capacities start at 1.13 bits and 1.06 at node A, for the low and high-noise cases, respectively. These are in good agreement with the predicted value 1.0 bits, showing that switchlike responses can accept reasonably large amounts of noise, and still respond nearly perfectly. The reason that the computed information is above the maximum theoretical information arises from the fact that the simulated systems are not ideal switches but have finite width transition zones (note that the green colors in Fig. 8H cluster around this transition zone). Moving downstream, the low-noise case preserves information almost perfectly, again showing that switchlike signaling is robust to small amounts of noise. However, information is rapidly lost in the high-noise case, showing its sensitivity to high amounts of noise.

## Discussion

This work investigates how dose-response curves between cellular signaling inputs and outputs can affect a signaling pathway’s ability to transmit information. It shows that pathways transmit more information when their dose-response curves have lower baselines, larger dynamic ranges, and curve shapes that correspond to the natural distribution of input signals. In particular, pathways can transmit more information if their dose-response curves are steepest around the input values that have the highest density of signals because this improves signal differentiation. If the signaling pathway is composed of multiple sequential steps, then the intermediate dose-response curves that represent the influence of one step on the next are called transfer functions. Here, we confirm prior suggestions that linear transfer functions typically improve information transmission. We also extend those results to show that closely related transfer function shapes can be equally good, and are in fact better in some cases (signaling pathways with multiple types of noise distributions). Additionally, we investigated switchlike transfer functions, finding that their signal transmission is very robust to small amounts of noise, but that they lose information rapidly when the noise that is high enough that there is a substantial probability of false-negative and false-positive responses. Stochastic simulations of toy signaling systems corroborate these findings.

An implicit assumption in this study is that biochemical reaction networks have functional modularity, meaning that their parts can be understood in isolation [92, 93]. In particular, we consider signaling pathways separate from the rest of the cell, and we also consider signaling pathways that are divided into sequential steps. This modularity assumption is essential for building a thorough understanding of how such systems work [94] and is supported by decades of research that has made the same assumptions and has come to sensible conclusions. Nevertheless, it’s worth noting that many signaling pathways interact with other pathways through crosstalk [95, 96] and that signaling pathways often include feedback loops [97, 98]. Additionally, there can be “hidden feedbacks” in a signaling pathway, or retroactivity, that arise from sequestration of upstream signaling molecules into downstream portions of the pathway [99]. Our work accounts for these interactions if they are internal to a single module, but not if they span multiple modules.

Our suggestion that dose-response curves can evolve to match natural signal distributions is not a new idea. As introduced above, Laughlin [33] showed that blowfly visual neuron contrast-response functions closely match the contrast levels in natural scenes, and he suggested that they evolved that way in order to optimize information transmission. It has also been suggested that gap gene expression levels in *Drosophila* embryos have been evolutionarily tuned to provide uniform precision for determining cellular positions along the anterior/posterior axis [100]. Although not recognized at the time as being important for information transfer, it has been observed that the binding constants of cell-surface hormone receptors tend to be reasonably close to circulating hormone concentrations [101]. This likely reflects the co-evolution of hormone production and receptor affinity to maximize the organism’s ability to respond appropriately to fluctuating hormone levels. Similarly, protein dimerization constants are often similar to their protein concentrations, allowing cells to use dimerization as a responsive regulatory switch rather than an obligate state [102, 103]. Both are consistent with dose-response curves evolving to improve signal transmission. As an aside, a concern in the field of biological information transmission has been that channel capacities may be poor measures of actual information transmission due to lack of knowledge of natural signal distributions [71]; however, this concern becomes moot if natural dose-response curves have evolved to match natural signal distributions because then cells do transmit information at the channel capacity.

For signaling over a single step, we showed that channel capacities are particularly sensitive to the bottom end of the response range for the gamma and constant-CV noise distributions. This suggests that cells would have evolved to minimize basal activation levels, such as by reducing protein autophosphorylation, leaky gene expression, and activation of receptors that don’t have ligands bound to them. Indeed, there is some evidence for this. Chapman and Asthagiri [104] found that overexpressing the yeast pheromone response protein Ste5 increased signaling pathway throughput but also increased the basal response. This suggests that Ste5 has a low natural concentration, despite the reduced maximal response and increased stochastic noise, in order to reduce the basal response and thus improve signal transmission.

We introduced the small and large noise approximations by stating that the SNA applies when the noise is much less than the response range and the LNA applies when the noise is similar to or greater than the response range. Our results suggest an alternate criterion. We found that graphs of channel capacity versus noise (right column of Fig. 3) are remarkably consistent across different noise distributions. The SNA channel capacity result is always within 20% of being accurate if it returns a channel capacity value of over 2 bits, while the LNA value is always within 20% if it returns a channel capacity below 0.9 bits (except for the constant-CV case, where the LNA approximation is poor, as discussed above). This shows that SNA results over 2 bits and LNA results under 0.9 bits are likely to be reasonably accurate, whereas in-between values require numerical calculation with the B-A algorithm.

The work described here is the final part of a larger investigation that focuses on three questions: (1) Do cells signal precisely? (2) If so, what approaches do they use? And (3) What biochemical mechanisms produce those approaches? We addressed the first question by showing that individual yeast cells do indeed signal precisely, transmitting over 3 bits of information in single static responses [10]. As mentioned in the introduction, this result is consistent with many other pieces of evidence [4–9]. The current work investigates the second question, showing that cells can tune their dose-response curves in order to better transmit information, including by aligning them with the natural signal distributions and by using linear transfer functions. These ideas had been suggested before [33, 43–47] but had not been shown rigorously. Finally, we addressed the third question of how cells can actually achieve linear transfer functions by systematically surveying toy signaling systems to see which ones can produce dose-response alignment [46]. We found that negative feedback with low gain is surprisingly ineffective, but that a push-pull mechanism is both effective and widely observed in cells [105–107]. Other mechanisms include negative feedback that’s coupled with high gain and protein activation from an effectively unlimited supply (we called this an “unsaturable cycle”), which are also observed in cells [41, 47]. Together, these projects present a picture of cell signaling systems that are highly evolved to transmit information precisely.

## Methods

We performed most of the work presented here with a combination of traditional paper-and-pencil mathematics and the Mathematica software. We drew reaction networks and conceptual diagrams with Microsoft PowerPoint and all other figures with Mathematica (except Fig. 4, which we created in Excel and PowerPoint). We did not use AI in any part of this work.

We performed all simulations described here with the Tellurium software [108, 109], which is a Python-based modeling tool that reads the human-readable Antimony language [110]. It performs stochastic simulations using the Gillespie algorithm, which is an event-based non-spatial simulation approach that has been shown to exactly represent its underlying chemical reaction model [111]. Sampling the simulation after every event introduces statistical artifacts, so we sampled it at regularly spaced time points instead. In preliminary work, we ran the simulations that are shown in Figures 8B and E in the Smoldyn software [112], which is a spatial stochastic simulator. The Tellurium and Smoldyn simulations agreed perfectly, within expected stochastic variation, further lending credibility to both simulators. See Supp. Inf. §7 for details.

## Appendix

Tkačik et al. claimed that it was optimal to set the LNA threshold at the crossing points of the two response distribution curves if they were unclipped Gaussians [69]. Here, we demonstrate that this is not the case.

The orange and green lines shown in Fig. 9 represent a pair of unclipped Gaussian conditional response distributions that are centered at 0 and 1, respectively. Taking their standard deviations *σ*_1_ and *σ*_2_, these functions are

**Fig 9.**
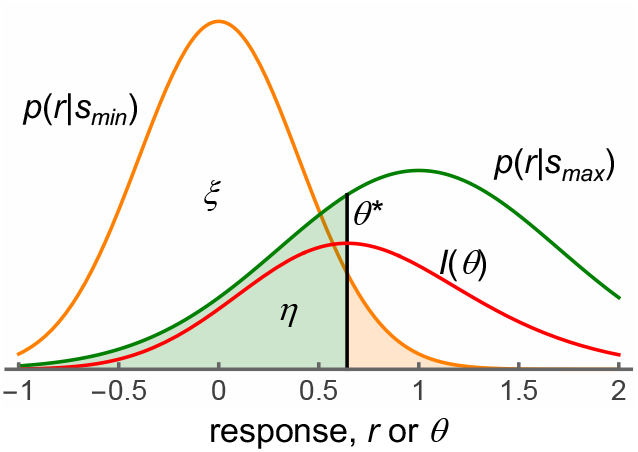
Threshold position for the LNA. The orange and green lines show unclipped Gaussian conditional response distributions that are centered at *r* = 0 and *r* = 1 and have standard deviations *σ*_1_ = 0.4 and *σ*_2_ = 0.7. Shading illustrates false-negative and false-positive probabilities. The red line shows the mutual information as a function of the threshold value. The peak of this red curve is the optimal threshold, which is around *θ*^*\**^ = 0.64. Notably, this optimal threshold is not where the orange and green lines cross, at about *r* = 0.51, and nor does it equalize shaded areas.

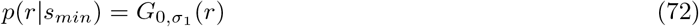

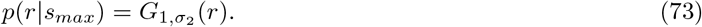

Substituting these probability distributions into Eqs. 14 and 15 yields the true-negative and false-negative probability values, each as a function of the threshold, *θ*,

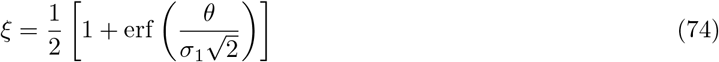

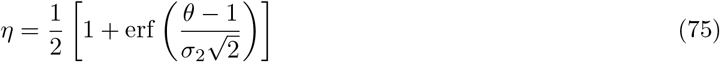

The red line in Fig. 9 represents the LNA mutual information result computed from Eq. 17, as a function of *θ*, using these probability values. The peak of this curve, illustrated with a vertical black line and labeled *θ*^*\**^, is the threshold value that maximizes the mutual information. It is not at the position where the conditional response distribution curves cross, although this may be an acceptable approximation in some cases. Perhaps more surprisingly, *θ*^*\**^ does not produce equal false-negative and false-positive probabilities, which are shown with shading (i.e. *η* ≠ 1*− ξ*). Instead, it appears that the optimal threshold can only be found numerically, as was done here.

## Supporting information

**S1 Supporting text and figures**.

## Author contributions

SSA conceived of the investigation, performed the research, and wrote the manuscript. RB had key initial insights. HMS and HSW helped edit the manuscript. All authors contributed to the discussion and acquisition of funding, and approved the final version of the manuscript.

## Acknowledgments

We thank Song Feng, Bill Peria, Lucian Smith, and Joe Hellerstein for helpful discussions. This work was supported by the National Cancer Institute under grant numbers U01CA227544 and R21 CA223901, the National Institute for General Medical Sciences R01GM155536-01, and a Simons Foundation Fellowship awarded to SSA. The content is solely the responsibility of the authors and does not necessarily represent the official views of the National Institutes of Health, the University of Washington or PNNL.

